# Planned missing data design: stronger inferences, increased research efficiency and improved animal welfare in ecology and evolution

**DOI:** 10.1101/247064

**Authors:** Daniel W.A. Noble, Shinichi Nakagawa

**Affiliations:** Ecology and Evolution Research Centre, School of Biological, Earth and Environmental Sciences, The University of New South Wales, Kensington NSW 2052, Sydney; Diabetes and Metabolism Division, Garvan Institute of Medical Research, 384 Victoria Street, Darlinghurst, Sydney, NSW 2010, Australia

**Keywords:** data augmentation, multiple imputation, personality, quantitative genetics, mixed effects models, hierarchical models, multilevel modelling, refinement, reduction, multiple working hypotheses

## Abstract

1. Ecological and evolutionary research questions are increasingly requiring the integration of research fields along with larger datasets to address fundamental local and global scale problems. Unfortunately, these agendas are often in conflict with limited funding and a need to balance animal welfare concerns.
2. Planned missing data design (PMDD), where data are randomly and deliberately missed during data collection, is a simple and effective strategy to working under greater research constraints while ensuring experiments have sufficient power to address fundamental research questions. Here, we review how PMDD can be incorporated into existing experimental designs by discussing alternative design approaches and evaluating how data imputation procedures work under PMDD situations.
3. Using realistic examples and simulations of multilevel data we show how a variety of research questions and data types, common in ecology and evolution, can be aided by using a PMDD with data imputation procedures. More specifically, we show how PMDD can improve statistical power in detecting effects of interest even with high levels (50%) of missing data and moderate sample sizes. We also provide examples of how PMDD can facilitate improved animal welfare and potentially alleviate research costs and constraints that would make endeavours for integrative research challenging.
4. Planned missing data designs are still in their infancy and we discuss some of the difficulties in their implementation and provide tentative solutions. Nonetheless, data imputation procedures are becoming more sophisticated and more easily implemented and it is likely that PMDD will be an effective and powerful tool for a wide range of experimental designs, data types and problems in ecology and evolution.

## Introduction

Missing data is a widespread problem in ecological and evolutionary research (Nakagawa & Freckleton 2010; Nakagawa & Freckleton 2011; Ellington *et al*. 2015; Nakagawa 2017), often resulting in the exclusion of a substantial amount of data. This contributes to a major reduction in statistical power and, if the nature of ‘missingness’ is not considered carefully, leads to biased parameter estimates (Enders 2001b; Graham 2009; Nakagawa & Freckleton 2010; Little *et al*. 2013). Theoretical frameworks for dealing with missing data, however, have received substantial attention and missing data theory is now a well-developed field of research grounded in solid statistical theory (Graham, Hofer & MacKinnon 1996; Enders 2001a; Little & Rubin 2002; Graham 2003; Graham 2009; van Buuren 2012; Little *et al*. 2013). Nonetheless, while social scientists have been at the forefront of applied missing data techniques, ecologists and evolutionary biologists have lagged behind (Nakagawa & Freckleton 2010).

Missing data is traditionally viewed by ecologists and evolutionary biologists with a sense of disdain and annoyance. But, what if including missing data in analyses could be advantageous? Indeed, social scientists have taken a rather different stance to missing data – instead embracing its power to help address fundamental research questions (Graham *et al*. 2006). Planned missing data design (PMDD) is an approach that involves deliberately planning to ‘miss’ data as an integral part of an experiment. In other words, deliberately not collecting data on certain variables or experimental subjects. While this seems like an odd thing to do, if missing data in the variables of interest is completely random, or can be made random, existing statistical frameworks are known to do an excellent job at estimating parameters and standard errors (Schafer & Graham 2002; van Buuren 2012). The PMDD approach comes with a substantial number of benefits that have been largely ignored by ecologists and evolutionary biologists.

Here, we argue that PMDD has the potential to expand the scope, reduce research costs and improve animal welfare, facilitating higher impact research with greater power. We begin our discussion by briefly introducing missing data theory and then describe a few core statistical tools that can be used to impute / augment (i.e., ‘fill in’) missing data. Using simulations, we show that, when missing data is ‘completely’ random, existing data imputation techniques can improve the estimation of parameters and their standard errors – even with hierarchically structured data that is common in ecological and evolutionary research (Enders, Mistler & Keller 2016; Quartagno & Carpenter 2016; Resche-Rigon & White 2016). We then describe PMDD, overviewing some of the different experimental approaches that can be implemented, what they involve and important design considerations. Following from this discussion, we overview the important benefits of utilizing a PMDD and end with a discussion on some of the challenges and limitations to their use – providing suggestions for how these can be rectified.

## A brief introduction to missing data theory

Missing data patterns can generally be classified as falling into one of three different types – based on the different mechanisms generating missing data – missing completely at random (MCAR), missing at random (MAR) and missing not at random (MNAR) (Rubin 1976; Little & Rubin 2002; Graham 2009; Nakagawa & Freckleton 2010; van Buuren 2012; Nakagawa 2017). The distinction between these three missing data mechanisms is important to understand when implementing a PMDD, which we will introduce below. Missing data (either in response or predictor variables) are considered to be MCAR when missingness is random with respect to both observed and unobserved (i.e., not collected in the study) variables (Enders 2001b; Nakagawa 2017). In other words, the observed data is simply a random sub-sample of complete data (Enders 2001b). In contrast, missing data are considered MAR when the missing values in a dataset depend on observed values of other variables in the dataset (Enders 2001b; Graham 2009). For example, if we were interested in understanding the correlation between survival to 1 year and mass at 6 months, we would find that individuals that die before 6 months are missing data on mass, but missing data on mass is correlated with their lifespan, which is known. Missing not at random (MNAR), however, occurs when missing values depend on unobserved variables that have not been quantified in the study, or on the variable itself after accounting for the observed data. For example, we may be missing behavioural data on small sized animals within a population because they tend to be ‘shy’ and difficult to capture (e.g, Biro & Dingemanse 2009), in which case we would be missing both behavioural and morphological data on a non-random sample of the population. Under these situations, dealing with missing data is difficult (possibly even impossible) because statistical techniques for estimating parameters when data are MNAR are difficult to implement given the need to explicitly model the process of missingness (Schafer & Graham 2002).

Missing data mechanisms have different consequences on statistical results when missing data is excluded prior to analysis, as is often the case (i.e., referred to as ‘complete case’, ‘pairwise deletion’ or ‘listwise deletion’). While MCAR results in a loss of power when missing data is excluded from an analysis, it does not bias parameter estimates (Enders 2001b; Schafer & Graham 2002; Graham 2009; Nakagawa & Freckleton 2010). In contrast, when missing data is MAR or MNAR, excluding missing data will result in both a loss of power and biased parameter estimates (sometimes severly so; Enders 2001b; Schafer & Graham 2002; Graham 2009; Nakagawa & Freckleton 2010). To better appreciate the impact missing data can have on sample size (and thus statistical power), assume that we have 10 variables, each containing 5% missing data, and a total complete dataset of *n* = 1000. If we used all variables in a statistical model we may need to exclude as much as 500 observations, resulting in a substantial decrease in power and severely compromising our ability to detect significant effects (see *‘Improving power of experimental designs’* for more on this issue). Statistical techniques for dealing with missing data rely on the assumption of missing data being MCAR or MAR, and if this assumption is met, then both power and bias in parameter estimates can be recovered (Enders 2001b; Schafer & Graham 2002; Nakagawa & Freckleton 2010; van Buuren 2012; Ellington *et al*. 2015; Nakagawa 2017).

## Statistical procedures for dealing with missing data

Planned missing data design hinges on the ability of researchers to make use of statistical procedures for handling missing data (Little & Rubin 2002; Graham *et al*. 2006; Enders 2010). It is therefore pertinent that we briefly review existing missing data procedures and provide some guidance on their implementation when data has been collected using a PMDD. We do not discuss these topics in great depth as there are a number of important, accessible reviews and books on these subjects already, which we direct the reader to for more details (Schafer 1997; Enders 2001b; Little & Rubin 2002; McKnight *et al*. 2007; Allison 2012; van Buuren 2012; Nakagawa 2017).

As mentioned above, imputation methods fall under two broad categories and we follow the general categorization of McKnight et al. (2007) in classifying them in to those implementing model-based (MB) techniques and those using multiple imputation (MI) with the help of Rubin’s rules (Rubin 1987; Enders 2010). While we acknowledge that these two broad categories have some overlap, they generally capture the major differences in the types of imputation procedures that can be applied. Model-based procedures incorporate both observed and missing data into a single joint modelling approach that is constrained by the underlying probability distribution and proceeds through the following steps: 1) the parameters of a model are estimated using observed data; 2) parameters estimated in step 1 are then used to augment missing data and 3) model parameters are re-assessed conditional on both the observed and imputed data (Figure 1A; Nakagawa 2017). These steps are reiterated until the model converges (i.e., maximum likelihood or stable posterior distribution if using Bayesian methods) (Figure 1A). Model-based approaches are advantageous in that they are fast, easily implemented (under the assumption of multivariate normality) and result in robust parameter estimates and standard errors (McKnight *et al*. 2007).

**Figure 1.**
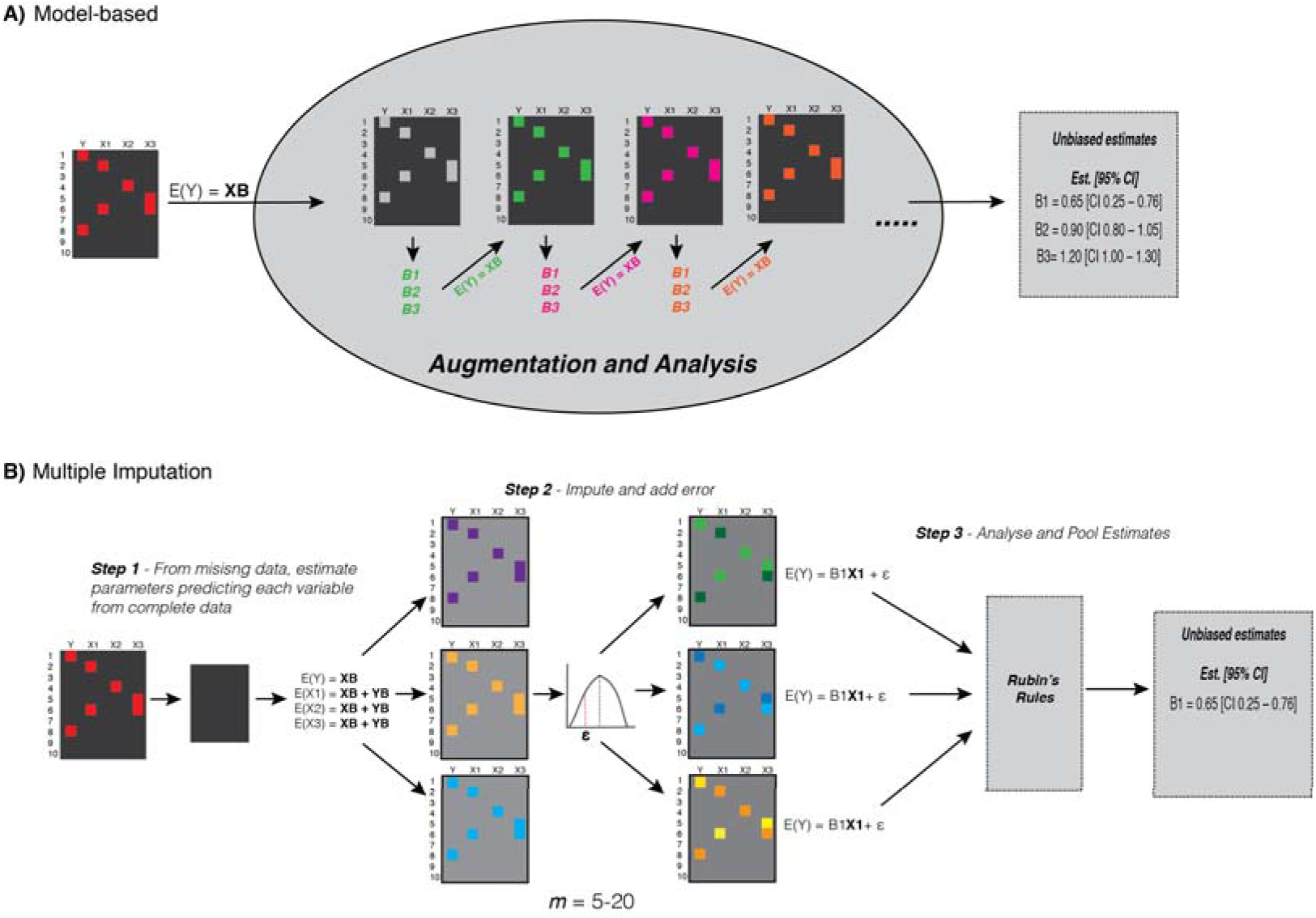
Two major types of imputation procedures: **A)** model-based procedures (e.g., full information maximum likelihood, expectation maximization, MCMC) and **B)** multiple imputation procedures. Each large square represents a dataset containing four variables (Y, X1, X2, X3) and n = 10 observations. Small red squares represent missing data and black squares complete data. Model-based procedures **(A)** take both observed and missing data in the analysis under a pre-specified model [E(Y) = **XB**)], augment missing data, estimate parameter estimates (B1, B2, B3) and then re-iterate this process with updated parameters [different coloured B1, B2, B3 and E(Y) = XB] until the model converges on a set of unbiased parameter estimates. Multiple imputation (B) uses complete data and often (but not always) uses other variables within the data as predictors of a specific variable. It then imputes using regression equations plausible values of missing data to generate *m* complete datasets. To prevent biased estimates residual error is added to each of the imputed data points (checkered small squares in step 2). These *m* datasets are then analysed with a given model, which can be different from the ones used to impute and, using Rubin’s rules (Rubin 1987), pool the parameter estimates across datasets (in this case B1). Abbreviations are as follows: E = expectation of variable, or mean estimate of variable; **X** = design matrix; B = vector of parameter estimates (e.g., B1, B2, B3); *ε* = residual effect or random error.

In contrast, multiple imputation proceeds by generating a set of *m* complete datasets where missing data is imputed using variables of interest. Usually *m* = 40–50 imputed datasets perform well under a variety of situations (Nakagawa & de Villemereuil 2015; Nakagawa 2017). These *m* datasets can then be analysed normally (i.e., as if a complete dataset existed) and the results (i.e., parameter estimates and standard errors) pooled across the *m* datasets (Figure 1B; Schafer 1997; Schafer & Olsen 1998; Little & Rubin 2002; van Buuren & Groothuis-Oudshoorn 2011; van Buuren 2012; Nakagawa 2017). Multiple imputation provides a number of advantages over MB approaches. First, it is extremely flexible, easily accommodating different distributions (i.e., Bernoulli, Poisson etc.), variables and model types if needed. Second, since MI creates *m* complete datasets it allows one to separate out the imputation step from the analysis step. In other words, we can have a set of auxiliary variables (see *‘Auxiliary variables to aid in imputation’* below) that are used to impute missing values and then subsequently use only the variables of biological interest to run the analysis on *m* imputed datasets. This is particularly advantageous because including unnecessary variables in MB procedures can complicate the interpretation of model results (McKnight *et al*. 2007; Enders 2010). Lastly, MI procedures account better for imputation uncertainty as variation in parameter estimates across data sets can be explicitly incorporated in pooled estimates, protecting against type I errors (McKnight *et al*. 2007). Additionally, the effect of missing data on analysis results (i.e. efficiency) can be explicitly quantified and presented by deriving statistics summarising the variability in parameter estimates across imputed datasets (McKnight *et al*. 2007). Given the flexibility, ease of implementation and their general tendency to produce robust parameter estimates, it is unsurprising that Rubin (1996) recommends MI over MB procedures.

### Auxiliary variables to aid imputation

Auxiliary variables are variables that are not necessarily of interest with respect to the biological question at hand, but that are correlated with other variables, or missing data itself, within the dataset (Collins, Schafer & Kam 2001; Graham 2003). Including auxiliary variables, especially ones that are expected to be predictive of missing data, have been shown to improve the accuracy and stability of estimates and to reduce their standard error (Enders 2010; Allison 2012; von Hippel & Lynch 2013). The best auxiliary variables are those that are easy and cheap to collect and that are strongly correlated with a number of other variables within the data set (Collins, Schafer & Kam 2001; Graham 2003; von Hippel & Lynch 2013). Collins et al. (2001) have shown that auxiliary variables can be particularly useful when the missing data is in the response variable, when they change the missing data mechanism from MNAR to MAR and when the correlation between auxiliary variables and the response is high (*r* = 0.9). Adding even just 2–3 auxiliary variables can improve imputation procedures and for the most part, an inclusive analysis strategy where a large number of auxiliary variables are included in the analysis is recommended (Enders 2010 p.g. 128). However, this procedure can be slightly more complex than this in practice. Hardt *et al*. (2012) show that the inclusion of too many (> 10) can start to lead to a downward bias in regression coefficients and a decrease in precision. In addition, auxiliary variables will have little impact when the correlations between variables in the dataset are low (*r* = 0.10) (Hardt, Herke & Leonhart 2012). While the number of auxiliary variables will depend on the specific study questions being asked, we recommend including auxiliary variables with moderate to high correlations (0.4–0.8). These will be particularly important when using imputation procedures where unplanned missing data exits (see section “*Unplanned missing data*” below) to change missingness from MNAR to MAR.

Experiments in ecology and evolution often collect variables that are not necessarily of interest, but can be used as auxiliary variables. These variables can include body dimensions, sex, age or even spatial data. These types of auxiliary variables can be included in imputation procedures (e.g., MI) with unplanned missing data to ensure that the MAR assumption is met, but then discarded when testing the biological questions and hypotheses of interest (Graham 2003). Considering these variables more carefully with respect to their potential correlations with other variables, and with missing data itself, is an important aspect of imputation because it can change missing data from MNAR to MAR satisfying assumptions of imputation procedures. As an illustrative example, consider a field study on birds, where the spatial position (i.e., latitude and longitude) of nest boxes is known and is stable through time. Here, the spatial position may not be of interest to researchers but it may be the case that it is correlated with behaviour (e.g., shyness) and / or body mass because subordinate animals get pushed to the fringes of habitat by dominant individuals and tend to be harder to recapture on repeated samples increasing the amount of missing data for these animals (Holtmann, Santos, Lara & Nakagawa, 2017). One way to use these spatial coordinates might be to generate a spatial covariance matrix between observations (using, for example, the SpatialTools package in R – French 2016) and extracting from it principle components (PCs). Multiple imputation could then make use of the PCs to impute missing data (e.g. using *mice* or *mi* – Table 3) for individuals that were not measured on a given sampling occasion. Similar approaches have been developed that make use of phylogenetic covariance matrices (Nakagawa & de Villemereuil 2015) as well as the relatedness matrices (Hadfield 2008).

## Planned missing data designs and their application in ecology and evolution

Planned missing data designs involve collecting incomplete data from subjects or observations of subjects on purpose by randomly assigning them to have missing measurements or measurement occasions (Graham *et al*. 2006; Rhemtulla & Little 2012; Little & Rhemtulla 2013). Researchers can then utilize model-based or multiple imputation techniques to fill in missing data such that the data contains complete information for all variables and experimental units within the dataset. Importantly, PMDD should always conform to the MCAR assumption because missing data is random by virtue of the experimental design making it ideal for use with imputation methods (Little & Rhemtulla 2013).

### Subset Measurement Design

Planned missing data design was first developed for research utilising questionnaires or surveys to help deal with participant fatigue, and is particularly useful when there are also logistical and financial constraints to asking many different questions (Graham, Hofer & Piccinin 1994; Graham *et al*. 2006). For example, a common type of PMDD called the *multiform design* (MFD) involves creating alternative questionnaires that each contain overlapping questions and a sample of new questions (Graham *et al*. 2006; Little & Rhemtulla 2013). Combining data on participants across the questionnaires, and then treating the questions participants were not given as missing information, allows missing data to be imputed based on the covariance between known answers (Graham *et al*. 2006).

In ecology and evolutionary biology, we often do not use questionnaires to collect data (aside from the field of ethnobiology; see Albuquerque *et al*. 2014), therefore, an analogous design is what we refer to as a *subset measurement design* (SMD) (Figure 2A). Similar to the MFD, a SMD involves quantifying a common set of variables across all individuals (e.g., body size) and then randomly allocating subjects to be quantified on a subset of other variables (e.g., hormone concentrations, metabolism etc.) (Figure 2A). Common variables can be those that are easily or cheaply quantified, such as body size indices (e.g., mass, body / wing length). In contrast, variables that are expensive or logistically challenging to quantify (e.g., gene expression, hormone concentrations) can be randomly sampled on a subset of subjects during the experiment. When using a SMD one should also consider, *a priori*, any potential interactions (Figure 2A) of interest and whether the planned missingness provides sufficient power to test these interactions (Enders 2010).

**Figure 2.**
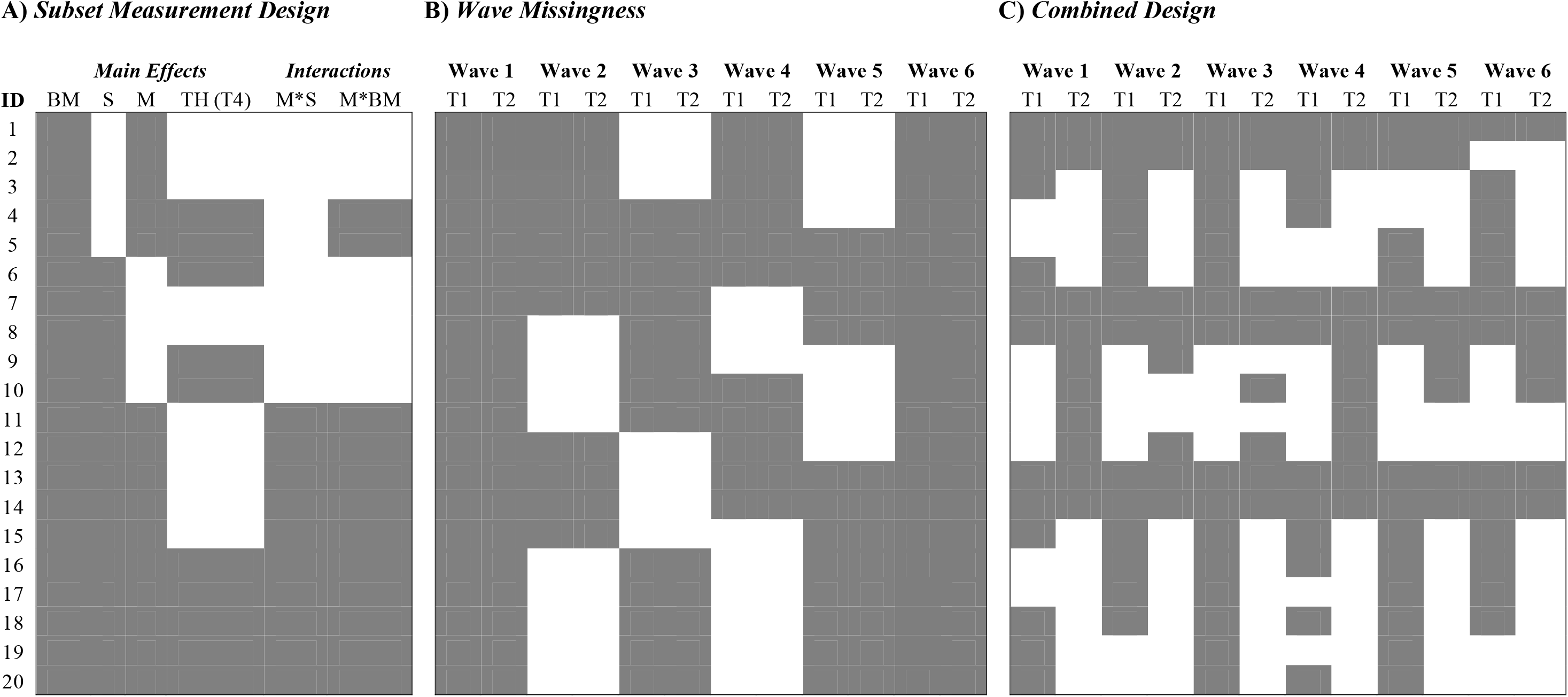
Three examples of planned missing data designs relevant for ecological and evolutionary research. In all cases, 20 individuals are shown along the rows (ID) and variables (e.g., BM) or traits (e.g., T1) measured are shown along columns. Abbreviations: BM = body mass, S = Sex, M = metabolism; TH = thyroxine (T4), T1 = Trait 1; T2 = Trait 2. ‘Gray’ filled boxes indicate that traits on individuals are measured and ‘white’ filled boxes are missed data. **A)** Subset measurement designs randomize a set of variables to be measured on a sample of individuals. Body mass (BM) is strongly correlated with all three other variables, and so, is measured on all individuals in the study. Molecular determination of sex is needed with our species, and along with thyroxine, can both be costly to quantify so these traits are measured on a sample of individuals. Metabolism can also be time consuming to measure, and so, it is only quantified on a sub-sample of animals. While we can estimate all main effects (i.e., single variables of interest) with this design, if interactions are of interest then one should have a PMDD that ensures there is enough data to effectively estimate the interaction parameters. **B)** Wave missing design can be applied to longitudinal data. Here, only two traits are quantified once a month on 20 individuals. In each measurement wave (i.e., a month), a random sample of individuals are completely lacking in data on both traits. **C)** Combined design. Longitudinal designs can miss entire sub-samples of individuals at lower rates, but also impose trait-level missingness for other individuals.

### Two-Method Design

SMD can also be applied to situations where researchers have a choice between two variables that might be related to some response variable of interest, but where one is more easily and cheaply quantified but has larger measurement error (i.e., less correlated with the response variable) and the second is more logistically challenging to measure but is considered the ‘gold standard’ (i.e., lower measurement error / more informative to the question). The latter design is referred to as a *two-method design* (TMD) in the social sciences (Little & Rhemtulla 2013). The two method design is often used in the social sciences when one variable is known to be systematically biased, which might be the case for self-reporting measures in surveys (Graham et al. 2006). Nonetheless, such approaches can still be used to help understand the relationship between a response variable and a more direct, but expensive measure, through the collection of a related, but cheaper and noisier variable. For example, we may be interested in measuring concentrations of thyroxine (T4) to understand how it relates to oxidative stress (concentrations of reactive oxygen species – ROS). Thyroxine is known to impact cell metabolism by being a involved in major hormone signalling pathways that affect ATP turnover in cells, and so, is expected to be related to ROS. However, T4 is costly to measure and we may be able to reduce costs by measuring a similar, more general ‘metabolic’ variable such as whole-organism resting metabolic rate, which is cheap and easy to measure but may not be as strong a predictor of ROS itself (e.g., Figure 2A – where M is measured more than T4). As such, depending on our question, we may actually measure more animals on whole-organism metabolic rate and fewer on T4, but still be able to understand the impact of T4 on oxidative stress itself.

### Wave Missingness Design

Longitudinal research questions, where repeated measurements on a set of independent individuals is of interest, can utilize a PMDD called *wave missingness* (Figure 2B). Here, a random group of experimental subjects are not measured at all at a given time point or measurement occasion (Little & Rhemtulla 2013; Rhemtulla *et al*. 2014). Waves can also be blocked such that all subjects are measured at the beginning and end measurement wave (e.g., month 1 and 6 – Figure 2B) with subsamples of animals not measured at all in middle waves (i.e., month 2–5 – Figure 2B) (i.e., pseudo-randomised missingness; Rhemtulla & Little 2012; Rhemtulla *et al*. 2014). Wave missingness designs have the potential to drastically decrease data collection costs because entire sets of individuals lack measurements at certain occasions. For example, if a study measured a set of traits on 20 animals across 20 populations, all populations might be sampled at months 1 and 6, but only some populations measured on months 2–5. Alternatively, we may be interested in understanding seasonal changes in profiles of two hormones across a sample individuals (e.g., testosterone and corticosterone), sampling the same set of individuals in a population at monthly intervals over the active season (6 months – Figure 2B). Here, we may take blood from all animals in months 1 and 6, but avoid sampling blood from a sub-sample of animals between months 2–5 Sampling animals across all sampling occasions means that we would need to sample blood 120 times and run 240 different assays. In contrast, with the missingness pattern in Figure 2B, we can reduce the total number of blood samples by ~20%, and avoid running 66 hormone assays (28% less). This can lead to reduced costs and stress to animals. The specific wave missingness design utilized will largely depend on the research question and the constraints faced in executing the study. However, careful attention needs to be paid when using wave missingness designs as they have been shown to perform poorly in certain situations with small sample sizes (Rhemtulla & Hancock, 2016).

### Considerations for and General Performance of Missing Data Designs

We have overviewed three of the more common designs that can be applied to experimental systems, however, it is important to note that PMDD’s can be diverse and are often not necessarily mutually exclusive of one another (Enders 2010; Rhemtulla & Little 2012; Little & Rhemtulla 2013; Rhemtulla *et al*. 2014). Combinations of the designs described above are probably necessary in many real research situations (e.g. Figure 2C). Regardless of which PMDD is used, researchers should choose the variables that are most pertinent to the specific hypothesis being tested, or those that are likely to have small effect sizes (and lower power), as those being measured with as little missing data as possible (i.e., having complete measurements on these variables). This ensures that the most pertinent questions can be tested rigorously (Graham *et al*. 2006). In addition, researchers should also consider the hypothesized correlation between variables. More tightly correlated variables (*r* > 0.50) may allow for one to plan for a greater level of missing data then two variables that are weakly correlated.

While PMDD’s can be powerful tools, it is still unclear what designs work best at estimating parameters and standard errors across various situations. Graham et al. (2006) and Enders (2010) (pg. 23–36) provide an excellent overview of the power of various PMDD’s. Graham et al. (2006) showed that with moderate sample sizes (*n* = 200–300), subset measurement type designs can have >0.80 power in estimating even small effects in many situations (i.e., Hedges’ *d* > 0.20). Rhemtulla et al. (2014) also show that with reasonably large sample sizes (*n* = 300 for multi-form design and *n* = 500 for wave missing and hybrid designs) that parameter estimates and standard errors in latent growth models show little bias. In many situations, any loss in power resulting from missing data seems to be rather small relative to the gains in the number of questions that can be tested and the logistical and cost improvements for a given experiment (Enders 2010). Despite simulations showing the benefits of PMDD, missing data itself can affect the information content in the data that is available for estimating parameters and standard errors (Rhemtulla & Hancock, 2016). It is important to recognize that missing information is not a simple function of the amount of missing data, it also depends on which variables are missing and the pattern of missingness (Rhemtulla & Hancock, 2016). Hence, a PMDD that is useful for, say, a linear latent growth curve model, may not be useful if the growth curves are non-linear. Indeed, efficiency – or the ability the to effectively estimate uncertainty in parameters – can be reduced when using PMDD relative to a complete case scenario (Rhemtulla et al. 2016).

## Benefits of planned missing data design

The PMDD’s outlined above provide a number of important advantages for ecologists and evolutionary biologists. Below we discuss these potential benefits more thoroughly using realistic simulations and examples to support our arguments. Despite the benefits, it is important to recognize, however, that PMDD’s do have limitations (see *“Challenges in implementing planned missing data designs*” section below), and despite their potential power in addressing a number of logistical constraints in research, it is unlikely they will be able to optimize all aspects of an experiment. When implementing PMDD it will be important to identify, *a priori*, what models will be fit to any data collected to make sure that the estimation of certain parameters can be done with high efficiency under the specific PMDD (Rhemtulla & Hancock, 2016).

### (i) Improving power of experimental designs

We have already indicated above that missing data procedures can improve the power of a given study by increasing the effective sample size. In the presence of missing data standard errors are estimated with less efficiency and thus the power to test the significance of an effect will decline (Little & Rhemtulla 2013). Studies in ecology and evolutionary biology are known to be under-powered in many cases (Møller & Jennions 2002; Jennions & Møller 2003), and so, this has important consequences for the inferences drawn in a given study. This is particularly true for multi-level data that often require large sample sizes to achieve sufficiently high power (van de Pol 2012) or even in genotype-phenotyping mapping studies, such as GWAS (here using imputation can also improve power; e.g., Marchini & Howie 2010). Integrating a PMDD into an experiment can recover the power lost after excluding missing data, facilitating the detection of small to moderate effects. To demonstrate how PMDD can improve inferences, even with multi-level hierarchical data, we conducted a couple simulations. In the first simulation, assume we are interested in estimating the between-individual level correlation between two traits (Y_1_ = “boldness” and Y_2_ = “time to emerge after predatory attack”) using a multi-response model. We simulated three variables (Y_1_, Y_2_ and Y_3_) that follow a multi-variate normal distribution (MVN) with between (B) and within (W)-individual covariance matrices (assuming a standard deviation (SD) = 1, which means the correlation and covariance matrices are the same) as follows:

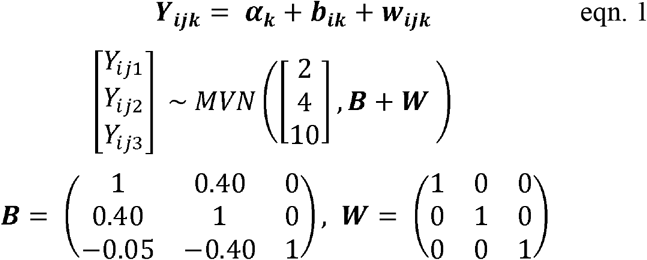

Here, is an **Y_*ijk*_** *j* x 3 matrix containing all observations, *j*, on each individual, *i*, for the three traits, *k* (Y_1_, Y_2_, and Y_3_); ***α_k_*** is a 1 x 3 scalar vector containing the intercept for each trait [2 4 10]*^T^* (means for Y_1_ – Y_3_); ***b_ik_*** is a *i* x 3 matrix containing the individual specific random effects or deviations for individual, *i*, for each trait, *k*, with 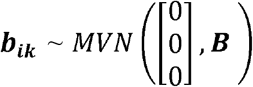. ***w_ijk_*** is the *j* observation level deviation for each individual *i* for the three traits *k* with 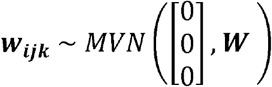. Note here that Y_3_ is not of interest with respect to the question, but because it is correlated with Y2 it can be used as an auxiliary variable to improve the imputation process. For this example, we simulated 5000 datasets under varying sample sizes. More specifically, the total number of observations ranged between 100 – 1000, with the number of individuals varying between 10–100, each being measured 10 times. The levels of missing data varied between 5–50%. We estimated the between– and within–individual covariance matrices under the above simulation scenarios using MB imputation (maximum likelihood approaches) in *ASReml–R* (vers. 3.4.1). Missing data was assumed to be MCAR throughout the data sets, as would be the case in a PMDD, and we evaluated how well imputation and complete case analyses performed in estimating the covariance between Y_2_ and Y_1_ under these varying situations (Figure 3). As expected, given our MCAR assumption, we did not see much impact on the point estimate (~0.40; Figure 3A & B). Imputation procedures actually tended to do slightly better at estimating the point estimate as there was slight estimation error for the complete case analysis with small sample sizes and high levels of missing data (Figure 3B). This was likely the result of model convergence problems when one has small sample sizes and high levels of missingness, with 15–26% of the simulations failing to converge under these conditions (i.e., *n* = 100 & 150 with 50% missingness). Despite this, we observed improvements in the estimation of standard errors when imputing data in comparison to the complete case analysis (Figure 2C & D), suggesting that imputation procedures, even with hierarchical data such as this, can lead to fairly substantial improvements in power. This is particularly important as many research areas, such as quantitative genetics and behavioural ecology, are interested variance partitioning methods to understand trait covariance (Dingemanse & Dochtermann 2013; Brommer & Class 2017; Careau & Wilson 2017).

**Figure 3.**
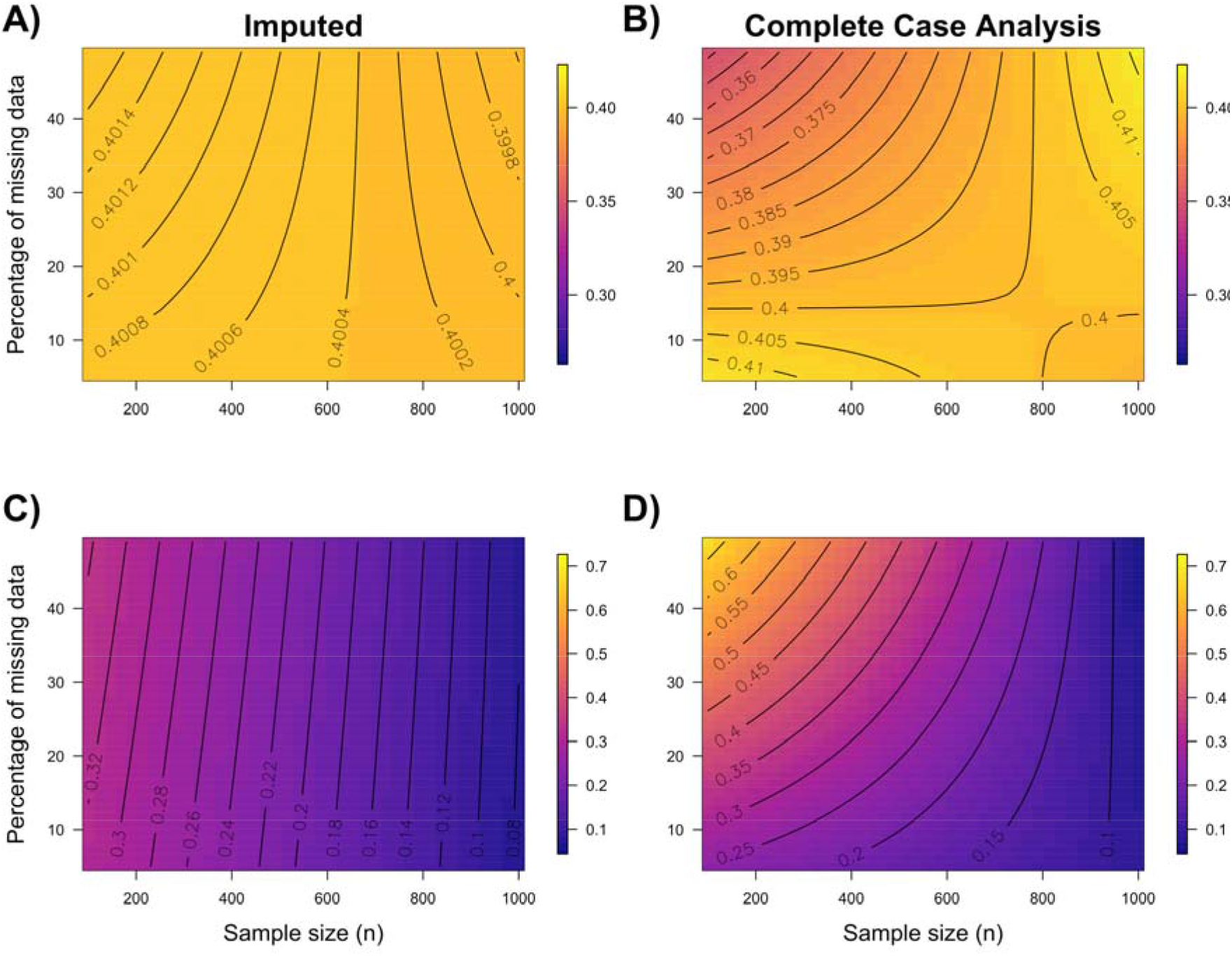
Simulation results of the estimation of a between–individual correlation for two traits [ (Y_1_, Y_2_) = 0.40] under varying levels of missing data (5–50%) and sample sizes (total number of observations; *n* = 100–1000, with number of individuals ranging from 10–100 each measured 10 times). Plots show average point estimates (A & B) and their corresponding standard errors (C & D) from 5000 randomly generated datasets when imputing missing data (A & C) or running a complete case analysis (excluding missing data – B & D). Note that convergence problems are more prevalent with small samples and high levels of missing data (see text). Parameter estimates and their precisions are therefore summarised on simulations in which models did converge.

Ecological and evolutionary questions are often more complex than simply estimating variance components. Experiments will often combine experimental manipulations of independent individuals and repeatedly measure these individuals across their life. To understand the benefits of PMDD at improving statistical inferences when both fixed and random effects might be of interest we conducted a second simulation. Assume we have manipulated the early thermal environment of a sample of lizard eggs – a common approach in lizard research (e.g., Noble, Stenhouse & Schwanz 2017). We might be interested in understanding how these thermal environments affect the growth curves of animals within each treatment (Figure 4). We simulated data assuming that individuals follow a linear growth trajectory (random regression model), at least over the period in which we measured their weight, according to the following model:

**Figure 4.**
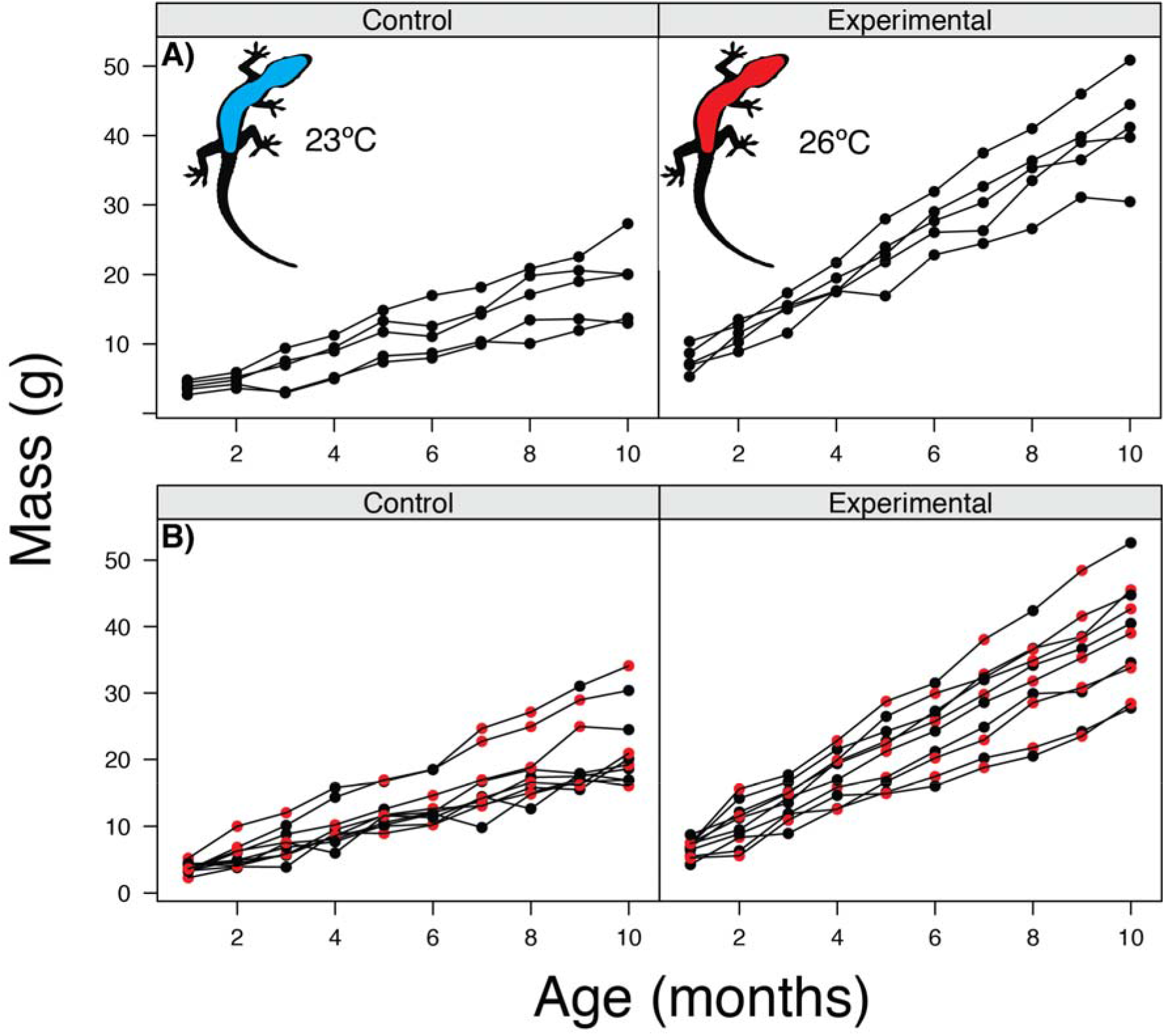
Example of experimental data on growth rates across age for two simulated scenarios. **A)** Mass of *n* = 10 lizards for a control group incubated at 23°C and experimental group incubated at 26°C. In this scenario, each animal was measured 10 times across the first 10 months of age. **B)** Mass of *n* = 20 lizards for a control group 23°C and experimental group incubated at 26°C measured five times with 50% of the mass data on each of the 20 animals considered missing (‘red’ points). In both scenarios, there were main effects of treatment and an interaction between growth across ages and treatment. Individual lizards varied in both their intercept and slope (see text for more details). Data was simulated according to eqn. 2 with the following parameters: *β*_0_ = 1.2; *β*_1_ = 2; *β*_2_ = 1.8; *β*_3_ = 1.6. In scenario B, missing data was imputed using likelihood based approaches in ASReml-R.

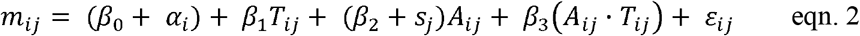

where *m_ij_* is the mass of individual *i* for observation *j*, *T_ij_* is a dummy variable (‘0’ or ‘1’) indicating whether observation *j* for individual *i* belongs to the control group (23°C) or the treatment group (26°C), *A_ij_* is the age of individual *i* at observation *j*, *β*_1_ is the contrast between the control group mean at age 0 (*β*_0_) and the treatment mean at age 0 *β*_2_, is the effect of age on mass, and *β*_3_ is the difference in mass change across age between treatments (i.e., interaction effect). *α_i_* and *s_j_* are individual level random effects for the random intercept and slope, respectively, which are assumed to follow an 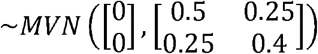. *ε_ij_* is the observation-level random effect assumed to follow an ~*N*(0, 1). An example of the simulated data is shown in Figure 4. If we were limited in the total amount of sampling, say 100 observations, then we can design an experiment where 10 animals (5 / treatment) were each measured 10 times (*Scenario 1*) or we could use a PMDD whereby we increase the total number of individuals to *n* = 20 (10 / treatment), but instead measure each individual randomly only five times, instead of 10 (*Scenario 2*) (Figure 4). Under these two scenarios we simulated 1500 datasets and used *ASReml-R* to estimate the parameters and their standard errors in eqn. 2. Overall, scenario 2 had a number of benefits both with respect to estimating fixed and random effects performing nearly as well as complete data (i.e., 20 individuals measured 10 times – *Scenario 3*). For all fixed effect estimates, there was between ~10–27% reduction in the standard errors with standard errors for the slopes and treatment interactions receiving a substantial boost. Interestingly, we also increase our precision in estimating random slopes (Table 1). Benefits of using a PMDD are even greater with larger sample sizes (e.g., 500 individuals measured 100 times *versus* 1000 individuals measured 50 times – see supplementary material Table S1), with a 23–29% reduction across both fixed and random effects. Estimation of random effects and their precision were improved quite a bit more with larger sample sizes (Table S1). Overall, our simulations suggest that PMDDs can provide power benefits under realistic experimental situations that are common in ecology and evolution.

**Table 1.** Effect of two scenarios on the ability to estimate parameters of a mixed model. Average parameter estimates and their standard errors for both fixed and random effects over 1500 simulated datasets. Example data is shown in Figure 4. Abbreviations and symbols as follows: *ρ*: correlation between intercept (*α*) and slope (*s*); 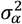 = variance estimate for random intercept; 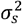 = variance estimate for random slope; trt = treatment; age = change in mass across age. % SE Decrease = the percent decrease in standard error from Scenario 1 compared to Scenario 2. “True” are the true parameters that the simulation was based on. Scenario 1 involves a hypothetical situation where 10 lizards are measured 10 times with 0% missing data. Scenario 2 is a situation where 20 lizards were measures 5 times (i.e., 50% missing data for each animal. In Scenario 3, 20 lizards are measured 10 times (i.e., a full complete case analysis for comparison). Note that simulations with larger sample sizes lead to even better improvements of SE when PMDD are implemented (Table S1).

|  | Scenario 1 |  | Scenario 2 |  | Scenario 3 |  | % SE<br>Decrease | True |
| --- | --- | --- | --- | --- | --- | --- | --- | --- |
| *Fixed Effects* |  |  |  |  |  |  |  |  |
|  | *Est.* | *SE* | *Est.* | *SE* | *Est.* | *SE* |  |  |
| intercept | 1.21 | 0.45 | 1.22 | 0.40 | 1.19 | 0.31 | 9.83 | 1.20 |
| trt | 2.01 | 0.63 | 1.98 | 0.57 | 2.00 | 0.44 | 9.82 | 2 |
| age | 1.79 | 0.28 | 1.80 | 0.21 | 1.79 | 0.20 | 27.06 | 1.8 |
| trt*age | 1.61 | 0.40 | 1.60 | 0.29 | 1.60 | 0.28 | 27.06 | 1.6 |
| *Random Effects* |  |  |  |  |  |  |  |  |
|  | *Est.* | *SE* | *Est.* | *SE* | *Est.* | *SE* |  |  |
| $\sigma^2_\alpha$ | 0.57 | 0.55 | 0.62 | 0.59 | 0.51 | 0.34 | -6.96 | 0.50 |
| $\rho(s, \alpha)$ | 0.24 | 0.25 | 0.23 | 0.21 | 0.24 | 0.16 | 16 | 0.25 |
| $\sigma^2_s$ | 0.41 | 0.21 | 0.41 | 0.15 | 0.40 | 0.14 | 31.26 | 0.40 |

### (ii) Improved animal welfare: considering the three ‘R s’

Central to research in ecology and evolutionary biology, particularly with respect to research on vertebrate animals, are issues surrounding animal welfare (Stamp Dawkins 2006; Barnard 2007; Cuthill 2007; Stamp Dawkins 2008; McMahon *et al*. 2012). Biological research on vertebrates, and indeed any study species, should strive to minimize pain, suffering and distress caused by experimentation, and we look to the three R’s (Refinement, Replacement and Reduction) to achieve these goals (Cuthill 2007). Planned missing data design can be an important tool during the experimental planning stages that directly targets two of the three R’s (Refinement and Reduction) to improve animal welfare. Often, PMDD can concurrently target both Refinement and Reduction at the same time by using less invasive procedures and allowing data to be collected on fewer experimental subjects (or less often on a given subject). For example, we could use a SMD to randomly assign the measurement of different physiological traits (e.g., metabolism, hormones) to a subset of subjects, reducing the number of subjects quantified on a given physiological measure. Additionally, fewer repeated physiological measurements can be done on a given subject by randomly assigning a different temporal sequence of measurements to individuals, refining the experimental design to reduce stress inflicted by repeated handling. Refinement can further occur by adopting a TMD where different physiological assays measuring the same construct, say ‘innate immunity’, can be done in such a way that no one individual has both measurements, but rather is randomly assigned to the cheaper less invasive measure. In summary, PMDD can improve animal welfare without compromising our ability to effectively answer a given question by designing an experiment that has too little power.

### (iii) Reduced research costs

One critical benefit of implementing PMDD is the ability to get a ‘bigger bang for your buck’ in terms of the research cost to outcome ratio. The cost savings when using a PMDD can be substantial, particularly for experiments involving expensive biochemical, proteomic, metabolomic and genomic work. However, they may also reduce costs for laborious experiments that require the breeding or measurement of a large number of individuals, such as is the case for diallel crosses in quantitative genetics (Lynch & Walsh, 1998). For example, assume we are interested in measuring hormone profiles for 60 individuals across a six-month activity period, meaning we would have a total of *n* = 360 blood samples (i.e., each animal sampled once a month). We might be able to reduce our sample size by 33% using a PMDD similar to that described in Figure 2C, bringing the total number of blood samples to *n* = 240. If we assume that the cost of individual reactions to run assays, in addition to labour costs, was $8 per sample, then we would save $900 for this experiment. These additional savings could be used to run a follow up experiment or even go towards assaying a sub-sample of individuals on a second hormone. Alternatively, we may also be interested in quantifying telomere length to understand cellular senescence using either flow cytometry (estimated cost / sample = $68) or qPCR (estimated cost / sample = $13) methods (Nussey *et al*. 2014) to understand patterns of telomere attrition over the season in relation to hormones. In this example, using a PMDD would save $8160 for flow cytometry methods or $1560 using qPCR methods (assuming we really had money to do all animals). It also demonstrates how a TMD can also save costs. Flow cytometry has been identified as a promising method for accurate, high throughput quantification of telomere length (Nussey *et al*. 2014). However, it is quite expensive compared to qPCR methods. A PMDD where a sample of animals are quantified both on qPCR and flow cytometry, and those missing flow cytometry data are then imputed would lead to substantial cost savings and allow one to verify the utility of both methods. We therefore view PMDD as a promising approach to improving cost efficiency.

### (iv) Stronger inferences and testing predictions from multiple working hypotheses

Strong inference involves devising alternative hypotheses and then running an experiment or set of experiments to test alternative predictions generated from these hypotheses (Platt 1964; Chamberlain 1965). Identifying and testing among alternative hypotheses is the hallmark of rapid scientific progress, indeed Platt (1964) suggested that this was one of the primary reasons for the rapid advancement of molecular biology through the 1960s. Nonetheless, it is clear that ecological and evolutionary studies rarely test predictions from multiple working hypotheses (Betini, Avgar & Fryxell 2017). Betini et al. (2017) suggest a number of intellectual and practical barriers impeding the use of multiple working hypotheses, but particularly relevant for our argument are the barriers limiting one’s ability to “execute” such investigations. Designing fully factorial experiments to disentangle predictions from alternative hypotheses is a major hurdle, referred to as the *“fallacy of the factorial design*” (Betini, Avgar & Fryxell 2017), whereby the addition of every new working hypothesis requires a new treatment or set of measurement variables leading to a geometric increase in number of required replicates. Including variables or experimental manipulations that test predictions from alternative hypotheses into a PMDD, can potentially off-set the need for prohibitively large sample sizes and time-consuming and costly measurements. These approaches may maximize the information gained from an experiment to facilitate more rapid scientific progress compared to testing single working hypotheses. Additionally, predictions from hypotheses can be tested using the same sample of experimental subjects, reducing the spatial and temporal differences between experiments.

### (v) Greater integration between research fields and disciplines

Understanding ecological and evolutionary processes require an integrative, multidisciplinary approach to tackling research questions (Wake 2009) – including the use of observational, experimental and theoretical modelling (Wake 2009). Integrative research questions cut across traditional boundaries, often making use of novel, and sometimes expensive techniques to help facilitate an understanding of a system or process. Despite its benefits, it can often be challenging to implement in practice given the different norms in various research fields, along with the costs and logistical constraints in doing integrative work. We argue that PMDD may help facilitate more multidisciplinary research efforts as it alleviates these constraints.

To demonstrate how PMDD can facilitate integrative research, assume that we are interested in testing Pace-of-life syndrome (POLS) theory (Reale *et al*. 2010). POLS theory explicitly predicts covariation between physiological, behavioural and life-history traits (Biro & Stamps 2008; Careau *et al*. 2010; Reale *et al*. 2010). Integration of suites of behavioural traits can lead to consistent individual differences in behaviour (i.e. personality – Reale *et al*. 2010; Stamps & Groothuis 2010) that can form behavioural syndromes (Sih & Bell 2008; Stamps & Groothuis 2010; Sih & Del Guidice 2012). Importantly, physiological mechanisms are thought to be the primary mechanisms (e.g. hormones, metabolism; Biro & Stamps 2008; Biro & Stamps 2010; Careau *et al*. 2010; Reale *et al*. 2010) underpinning both behavioural and life-history variation in populations. Pace-of-life theory is integrative, requiring concurrent measurements of physiological, behavioural and life-history traits to understand their covariance. It often requires laborious, time intensive, and sometimes costly measurements of the same individuals over time, posing major hurdles in obtaining the necessary data required. Additionally, physiological measures (e.g. metabolism, hormones, ROS, immunity) can be costly to obtain limiting the number of samples that can be collected. Despite this, it is important to take many physiological measurements (Adamo 2004). A PMDD may alleviate some of these constraints by reducing the need to measure all variables across subjects over time.

## Challenges in implementing planned missing data designs

As with any new research method, there will be challenges, particularly in establishing the most suitable approaches that work across a wide diversity of different research questions and experimental designs in ecology and evolution. Given that PMDD is still new, stimulating interest in them will be the first step to identifying, solving and implementing solutions to some of the challenges that crop up. Below we discuss some of the hurdles we see to implementing a PMDD and suggest some tentative solutions.

### Unplanned missing data

Of course, as with any experiment, unplanned missing data will creep into PMDD designs. Often missing data may be random, such as when a piece of equipment malfunctions during a set of measurements or when recording errors are identified. Random instances of missing data, even if unplanned, will not affect the imputation process or the utility of PMDD unless missing data levels begin to get quite high. However, our simulations show, as well as others’, that imputation procedures perform quite well even with large amounts of missing data (~50% – see Figure 3). Nonetheless, there are real situations where unplanned missing data can be MNAR and this will affect any experiment regardless of whether a PMDD is implemented or not. We have outlined above how data can be made MAR though the use of auxiliary variables, and these unplanned missing data, can then be imputed normally along with any planned missing data using the same statistical methods. We therefore advise colleagues to collect possible auxiliary variables where possible to counter unplanned missing data.

### Imputation with generalised linear mixed effect models

Multiple and model-based (MB) imputation both work well with normally distributed data, however, in reality variables often are non-normally distributed. While MB procedures are limited to situations that assume multivariate normality, MI procedures can also work with non-normal data fit using generalised linear mixed effect models (e.g., Poisson GLMMs) (Schafer 1997). However, implementation in the context of GLMMs is still under active development, and in many cases, is restricted to simple random effect structures (van Buuren & Groothuis-Oudshoorn 2011; Enders, Mistler & Keller 2016; Quartagno & Carpenter 2016; Audigier & Resche-Rigon 2017). Nonetheless, two-level random regression models can be run in a number of existing packages (e.g., *mice*) and we believe that the capacity to run more sophisticated models will grow in the near future.

To re-assure readers that imputation can and does work with non-normal distributions, we provide a simulated example along with sample R code to demonstrate MI procedures with GLMMs in Box 1. For our hypothetical example, assume we are interested in provisioning rates (the number of feeding visits by a parent) in a bird species, the fictitious Missing Capped Warbler (*Sylvia absenscapilla*). We would like to understand the costs of female provisioning by experimentally manipulating brood sizes (*n* = 6 chicks) in a random sample of birds compared to a control group, which has normal brood sizes (*n* = 3 chicks) (Liebl, Browning & Russell 2016). The Missing Capped Warbler is notoriously difficult to observe as it is found in thick scrub, and so, we placed cameras at random nests during the first two weeks of the breeding season to observe provisioning rates in the two treatments over a 5-hour period. Provisioning rates are known to change as chicks develop (Khwaja *et al*. 2017), and so, the cameras were on each nest for a total of 20 days to understand how the demands of chicks change, and whether females can keep up with these demands. Unfortunately, it is a laborious process to observe all the resulting video for 40 birds measured over 20 days (a total of 4000 hours of video!). We therefore decided to implement a PMDD, where we randomly sampled a set of videos (n = 536 out of 800 videos; ~ 33% missing data) where provisioning rates can be quantified (cutting the total hours of video watching to 2680 hours). Planned missing data can be imputed, for example, using the *mice* package (see Box 1). After a long and laborious field season, we were able to collect data that shows a tendency for experimentally elevated clutch sizes to have higher rates of provisioning that increase over the care period (Figure 5). Overall, we see that the imputed data matches well with the complete dataset, with nearly identical results in this simulation (see Table in Box 1).

**Figure 5.**
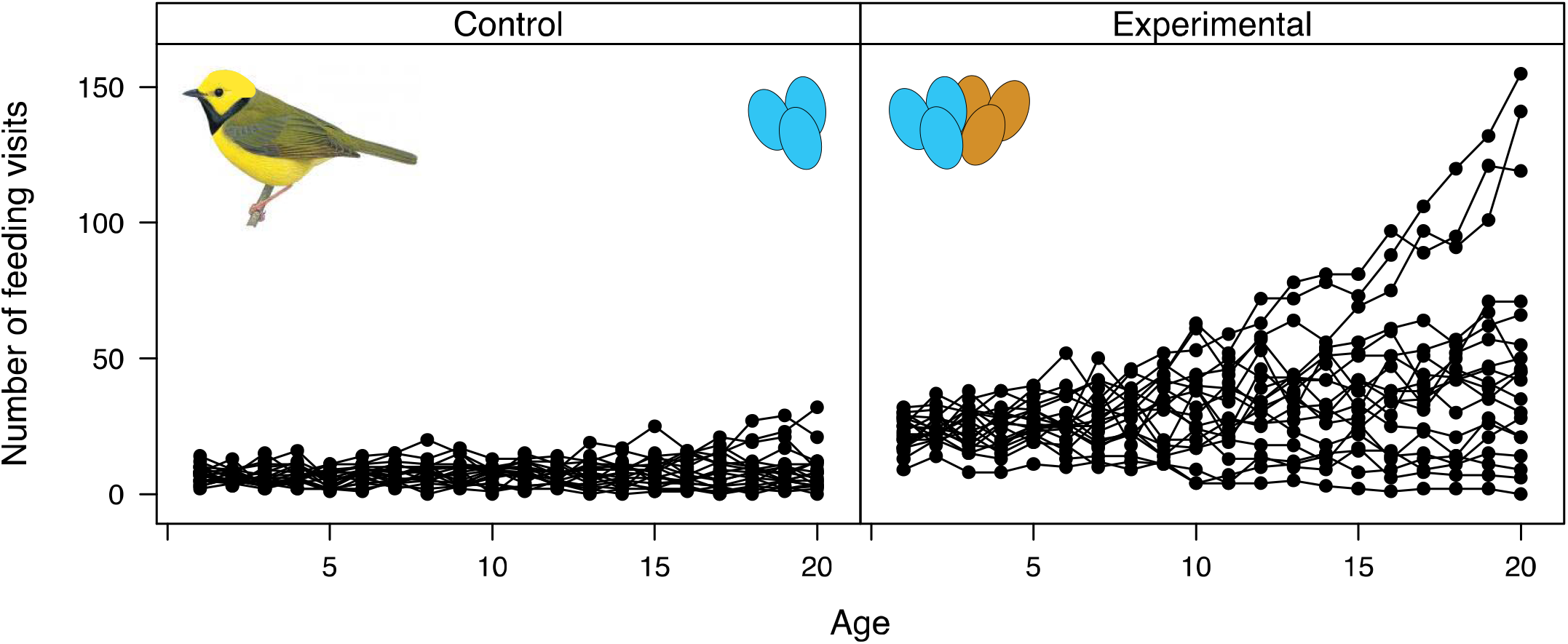
Example data showing the provisioning rates (number of feeding visits within a 5-hour period) for control (3 eggs) and experimentally elevated (6 eggs) brood sizes in the fictitious Missing Capped Warbler across the first 20 days of chick age. Provisions were simulated assuming a Poisson error distribution using the following model: *n_ij_* = (*β*_0_ + *α_i_*) + *β*_1_*T_ij_* + (*β*_2_ + *s_j_*)*A_ij_* + *ε_ij_* where Provisions = log(*n_ij_*). *β*_0_ = 2; *β*_1_ = 1; *β*_2_ = 0.01 with a random intercept and slope variance and covariance matrix as: 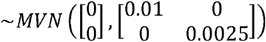.

### Overcoming psychological barriers to missing data

One of the biggest challenges to implementing PMDDs probably involves the need for researchers to over-come the ‘psychological taboos’ around missing data, and the suspicion of techniques for handling missing data (Enders 2010). We can re-assure readers that missing data practices are now very well established (Graham *et al*. 2006; Nakagawa 2017), and are rather painlessly implemented in many commonly used statistical software such as *R*, *SAS*, *SPSS*, and *MPlus* (See Table 2 for an overview). In fact, many techniques are implemented by default when missing data is included as response variables in models for a number of mixed modelling packages (e.g. model-based procedures in ‘*MCMCglmml* and ‘*ASReml*). While statistical algorithms vary across these platforms, fairly sophisticated and versatile ones are now implemented in packages for some of the most widely used platforms (e.g. *‘mice’* ‘*mi*’, *‘multimput’* and ‘*Amelia*’ in the R environment – Table 2) that implement MI algorithms known to perform well under a wide variety of situations (Schafer & Graham 2002; van Buuren 2012; Enders, Mistler & Keller 2016; Quartagno & Carpenter 2016; Resche-Rigon & White 2016; Audigier *et al*. 2017). These techniques are under active development (e.g., the *mice* package in R), and so, we envisage the breadth of problems these tools can tackle to increase and be even easier to apply in the future. Nonetheless, caution is still needed in their implementation as it is unclear whether imputation procedures perform well under all circumstances (Nakagawa 2017). Statistical procedures for missing data are still rarely taught in undergraduate and graduate level courses, so part of the solution will be to begin educating students and practitioners about how to perform imputation procedures, explicitly highlighting some of the challenges and caveats that need to be considered. Nonetheless, there are now excellent resources that provide an in depth look at imputation procedures (Gelman & Hill 2002; Schafer & Graham 2002; Graham 2009; Enders 2010; Su *et al*. 2011; van Buuren & Groothuis-Oudshoorn 2011; Nakagawa 2017).

**Table 2.**
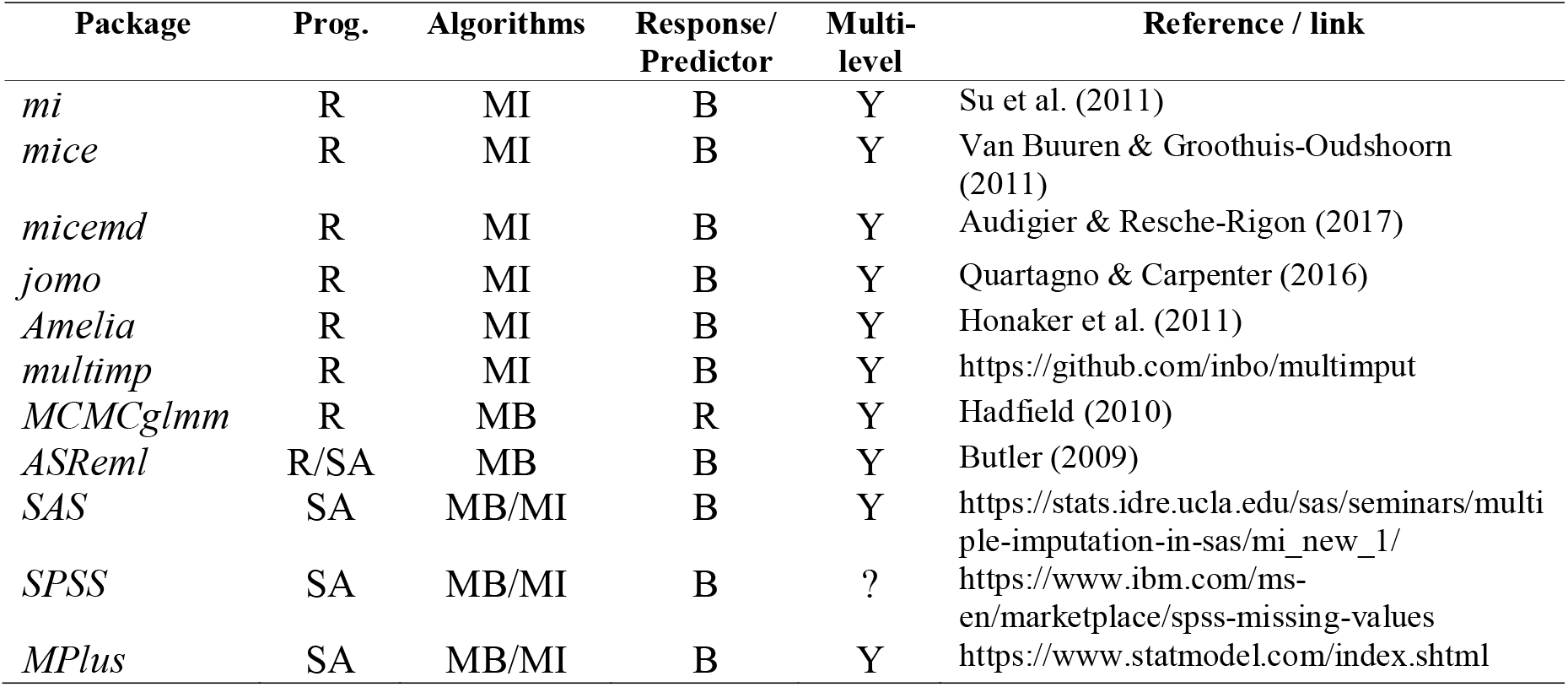
Examples of common packages and statistical programs that can be used to deal with missing data. Abbreviations are as follows: SA = stand-alone program; MI = multiple imputation; MB = model-based imputation; B = both, R = response, P = predictors, Y = yes, ? = unknown.

| Package | Prog. | Algorithms | Response/<br>Predictor | Multi-level | Reference / link |
| --- | --- | --- | --- | --- | --- |
| *mi* | R | MI | B | Y | Su et al. (2011) |
| *mice* | R | MI | B | Y | Van Buuren & Groothuis-Oudshoorn (2011) |
| *micemd* | R | MI | B | Y | Audigier & Resche-Rigon (2017) |
| *jomo* | R | MI | B | Y | Quartagno & Carpenter (2016) |
| *Amelia* | R | MI | B | Y | Honaker et al. (2011) |
| *multimp* | R | MI | B | Y | https://github.com/inbo/multimput |
| *MCMCglmm* | R | MB | R | Y | Hadfield (2010) |
| *ASReml* | R/SA | MB | B | Y | Butler (2009) |
| *SAS* | SA | MB/MI | B | Y | https://stats.idre.ucla.edu/sas/seminars/multiple-imputation-in-sas/mi_new_1/ |
| *SPSS* | SA | MB/MI | B | ? | https://www.ibm.com/ms-en/marketplace/spss-missing-values |
| *MPlus* | SA | MB/MI | B | Y | https://www.statmodel.com/index.shtml |

### Uncertainties surrounding the best PMDD

One challenge in implementing PMDD is the uncertainty around what the most appropriate missing data design is for a given experiment. This is particularly true in ecology and evolutionary biology because different questions, experimental systems, data structure and measurement variables may require creative combinations of different PMDD’s that we discuss in our paper – or possibly even new ones! While we argue that the benefits of PMDD can be substantial it may still be important to test the robustness and power of any given PMDD through simulations (Rhemtulla & Handcock, 2016). With some very simple simulated data based on effect sizes and experimental designs relevant to the question at hand, the power of different PMDD’s can be thoroughly tested during the design stage of an experiment (Enders 2010). This may require researchers thinking carefully about the model they wish to test and simulating data under this model early on (Rhemtulla & Handcock, 2016). Enders (2010 p.g. 30) provide a nice introduction on how to conduct power analysis with PMDD’s using simulations, and we provide all our R code which we hope can act as a skeleton for readers to familiarize themselves with simulations to help them sort out the best PMDD for their particular situation. Additionally, new multi-level simulation packages, such as SQuID, allow for researchers to simulate hierarchical data easily (Allegue *et al*. 2017). Data can be downloaded and missing data introduced to evaluate the power of different PMDD’s. While definitive guidelines will depend on the experimental design, question and covariance between traits, we believe that up to ~30% missing data overall across a wide variety of different designs will likely not compromise performance of imputation procedures.

## Conclusions and future directions

Our goal was to put planned missing data design on the radar of ecologists and evolutionary biologists given the substantial number of ethical, logistical and cost saving benefits it may afford. We have provided some guidance on possible PMDD’s that can be implemented in research programs and shown, with simulations, that even hierarchical / multilevel data imputation procedures can perform quite well. While it is still unclear whether imputation procedures in a multilevel framework will work under all circumstances there is increasing awareness of the need to develop such techniques with hierarchical data and in many cases existing methods will likely perform well (Drechsler 2015; Quartagno & Carpenter 2016; Audigier & Resche-Rigon 2017; Audigier *et al*. 2017). We encourage colleagues to begin thinking about PMDD’s and their utility in their research both to improve research quality and to promote integrative, cost effective research projects in ecology and evolutionary biology.

## Acknowledgements

We would like to thank the I-DEEL lab (Joel Pick, Rose O’Dea, Malgorzata Lagisz, and Fonti Kar), Diego Barneche and Corey Callaghan for feedback on previous versions of our manuscript. We would also like to thank the editors, Mijke Rhemtulla and an anonymous reviewer for very constructive comments that greatly improved our manuscript. DWAN was supported by an ARC Discovery Early Career Research Award (DE150101774) and SN an ARC Future Fellowship (FT130100268). The authors declare no conflicts of interest.

## Author’s contributions

SN and DN conceived the research. DN and SN conducted the simulations. DN wrote the manuscript, with input from SN. SN contributed critically and constructively to the revisions of the manuscript.

## Data Accessibility

All R code and simulation results used in the manuscript were provided upon submission.

### BOX 1

Multiple imputation (MI) for a few different generalised family types can be implemented in the *mice* package (van Buuren & Groothuis-Oudshoorn 2011). Given that our data (number of visits in 5 hours) is Poisson distributed (or nearly so), and hierarchical in nature, we will need to impute using a new addon developed in the *countimp* package (Kleinke & Reinecke 2013), which imputes Poisson random regressions. It can be installed, along with other needed packages in R as follows:

~~~
> link <- “http://www.uni-bielefeld.de/soz/kds/software/countimp_1.0.tar.gz”
> install.packages(link, repos=NULL, type=“source”)
> library(countimp)
> library(mice)
> library(glmmADMB)
~~~

Our data, including the missing data is set up as follows:

~~~
> head(data)
 ind age trt trt_name provision
1 1 1 1 Experimental NA
2 1 2 1 Experimental 22
3 1 3 1 Experimental 29
4 1 4 1 Experimental 38
5 1 5 1 Experimental 33
6 1 6 1 Experimental 37
~~~

In total, we have approximately 33% missing data at the individual level. To impute these missing data, we need to first set up the predictor matrix to define what variables are class (random effect groups) and what are to be used as random and fixed effects:

~~~
> data$ind <- as.integer(data$ind) # Need to keep class variable as integer
> imp <- mice(data, maxint = 0, printFlag = FALSE) *#* Do quick run of mice to set up pred matrix
> pred <- imp$pred # Extract the pred matrix
~~~

Now that we have the predictor matrix, to run a multi-level imputation we need to change the predictor matrix row for provision to define the class variable (i.e., random effect group – set as ‘−2’ – only one level can be included currently), the variable included as both a fixed and random effect (i.e., age – set as ‘2’ because we have a random regression model) and we will set our ‘tr? as a fixed effect only (i.e., set as ‘1’) as follows:

~~~
> pred[“provision”, ] <- c(−2,2,1,0,0)
~~~

The ‘0’ is used to tell mice not to include these variables as predictors in the imputation. Now that this is set up we can run multiple imputation telling mice that the ‘provision’ variable is a 2-level Poisson variable:

~~~
> imp <- mice(data, m = 20, meth = c(““,”“, “21. poisson”), pred = pred)
~~~

This will impute missing information in the ‘provision’ variable creating *m* = 20 ‘filled in’ datasets for which estimates and standard errors can be pooled as follows:

~~~
> fit <- do.mira(imp = imp, DV = “provision”, fixedeff = “trt+age”, randeff = “1 + age”, grp = “ind”, id = “ind”, fam = “poisson”)
> summary(fit)
~~~

We can compare the overall pooled estimates with estimates from our ‘real complete dataset’ (assuming we watched all videos):

| Full Dataset |  |  | Imputed Dataset |  |
| --- | --- | --- | --- | --- |
| Fixed Effects |  |  | Pooled Fixed Effects |  |
|  | Estimate | SE | Estimate | SE |
| Intercept | 1.93 | 0.05 | 1.93 | 0.06 |
| trt | 1.13 | 0.06 | 1.13 | 0.07 |
| age | 0.01 | 0.01 | 0.01 | 0.01 |
| Random Effects |  |  | Pooled Random Effects |  |
|  | Estimate |  | Estimate |  |
| Intercept | 0.011 |  | 0.015 |  |
| Age | 0.0025 |  | 0.0023 |  |

